# Does the margin of stability measure predict stability of gait with a constrained base of support?

**DOI:** 10.1101/646943

**Authors:** Lakshdeep Gill, Andrew H. Huntley, Avril Mansfield

## Abstract

This study aimed to determine the validity of the centre of mass position (COM) position and extrapolated COM (XCOM), relative to the base of support, for predicting stability during a walking task where the base of support is constrained. Nine young healthy participants walked on a narrow beam. Three-dimensional motion capture was used to calculate the COM and XCOM relative to the base of support. Steps were classified as having either the COM or XCOM inside or outside the base of support, and were classified as successful (stable – foot placed on the beam) or failed (unstable – foot stepped off the beam). If the COM or XCOM are valid measures of stability, they should be within the base of support for successful steps and outside the base of support for failed steps. Classifying the COM and XCOM inside or outside the base of support correctly predicted successful or failed steps in 69% and 58% of cases, respectively. When the COM or XCOM were outside the base of support, walking faster seemed to help participants to maintain stability. The further the COM or XCOM were outside the base of support during a successful step, the more likely participants were to fail on a subsequent step. The results of this study suggest that both COM and XCOM are valid measures of stability during a beam walking task, but that classifying COM and XCOM as inside or outside the base of support may be over-simplistic.

## INTRODUCTION

Mechanical stability of a system can be described by the relationship between the centre of mass (COM) and the boundaries of the base of support (BOS; Pai and Patton, 1997; Hof *et al.*, 2005). Traditionally, the system is considered ‘stable’ if the COM is located within the BOS (Shumway-Cook and Woollacott, 2007); however, this assumption fails during movement (e.g., normal overground bipedal walking), when the COM is often located outside of the BOS (Winter, 1995). The margin of stability (MOS) has been proposed to quantify mechanical stability in complex, dynamic situations where balance is maintained even if the COM is outside the BOS (Hof *et al.*, 2005). MOS is the difference between the position of the extrapolated COM and the edge of the BOS (Hof *et al.*, 2005; Hof, 2008); where the extrapolated COM considers the velocity of the COM in addition to its position in space. MOS indicates how close an inverted pendulum is to falling; a positive MOS (extrapolated COM inside the BOS) suggests mechanical stability whereas a negative MOS suggests instability (Hof *et al.*, 2005; Hof, 2008; Bruijn *et al.*, 2013). In cases of mechanical instability, corrective movements such as changing step placement to increase the size of the BOS would be required to prevent a fall (Hof *et al.*, 2005; Hof, 2008).

Despite the utility of MOS for describing mechanical stability during human movement, MOS calculations are problematic under certain circumstances. For example, during the single support phase of normal bipedal walking, MOS will be negative relative to the anterior and medial boundaries of the BOS (Bruijn *et al.*, 2013; Sivakumaran *et al.*, 2018). However, the system is not strictly ‘unstable’ in these circumstances as the next planned step will be placed anterior and medial to the stance limb. This calls into question whether the MOS can be applied to all phases of the gait cycle.

This study aimed to assess the validity of COM position and MOS describing medio-lateral mechanical stability during gait when the width of the BOS is constrained; restricting the width of the BOS prevents participants from compensating for instability by adjusting step placement (Bruijn and van Dieën, 2018; Sivakumaran *et al.*, 2018). Participants walked on a narrow balance beam, such that the width of the beam is the width of the BOS. To perform this task correctly the swing foot must be placed in tandem on the beam in front of the stance foot, and instability is clearly observed as a step off the beam and a ‘failure’ of the task. We hypothesized that the MOS would more accurately describe mechanical stability than the COM position relative to the BOS. The accuracy of each measure in describing mechanical stability was defined as the proportion of correctly predicted successful and failed steps.

## METHODS

### Participants

Data were collected from nine healthy young adults (20-35 years old) with no neurological or musculoskeletal conditions limiting mobility. Participants were excluded if they had more than three months of training in dance or gymnastics at any point in their lives. Participants were six women and three men, with a mean (standard deviation) age, height and weight of 23.3 (3.2) years, 164.5 (11.7) cm, and 65.3 (12.6) kg, respectively. Participants provided written informed consent prior to participation. The study was approved by the University Health Network Research Ethics Board.

### Procedure

Prior to the beam-walking trials, participants performed three 4-m tandem walking trials along a flat walkway. The number of steps and the amount of time taken to complete the walkway were recorded; number of steps were divided by time to calculate cadence; this cadence was used to set a target cadence for the beam-walking trials (see below).

Participants were outfitted with 58 reflective markers with marker layout according to Visual3D marker set guidelines (C-Motion, Germantown, USA) guidelines. Marker clusters were used on the chest, pelvis, thighs and shanks. Individual markers were placed on the head, acromion processes, pelvis, knees, ankles and shoes. Kinematic data were collected using a Vicon Mx 14-camera system (Vicon Motion Systems Ltd, Oxford, UK). Participants performed four beam-walking trials on a wooden beam (4 m long, 3.8 cm wide, and 8.5 cm high) bolted onto the floor. The starting position along the beam was adjusted for each participant depending on their shoe length, which ensured that, if each trial was executed perfectly (participant stepped directly heel-to-toe with no space between the feet), the participant would take ten steps per trial. Participants were instructed to walk with their steps in heel-to-toe fashion and arms folded across their torso, while matching their footfalls to the beat of a metronome. The metronome rate was set at 80% of participants’ over-ground cadence; a slower cadence was used as walking on the beam is more difficult than tandem walking on a flat surface. Participants were instructed to land their first footfall on the 5^th^ beat of the metronome, and to land each subsequent step on the next beat. Participants were also told not to look at their feet, and to keep their eyes directed ahead to a point on the wall. To ensure that participants could not see their feet participants wore basketball ‘dribble goggles’ that obscured the bottom half of the visual field. If the participant lost balance and stepped off the beam, s/he was instructed to step back at the same point on the beam. The trial ended when the participant reached the end of the beam. Participants wore a safety harness attached to a secure overhead track during all trials to prevent a fall to the floor. All trials were video recorded using four cameras placed around the laboratory.

### Data processing

Motion capture data were filtered in Visual3D using a second order, low pass Butterworth filter with a cut-off frequency of 6 Hz, and marker gaps of <10 frames were interpolated. Foot strike was defined as the foot touching either the beam or the floor, which was identified as the time when the antero-posterior velocity of the heel marker was <0.1 m/s (Ghoussayni *et al.*, 2004), and confirmed with visual inspection. Each step was defined from foot strike to foot strike; for example, steps with the right foot were from left foot strike to right foot strike. The success of each step was coded from the video footage. Successful steps were steps where the swing limb landed on the balance beam and failed steps were steps where the swing limb landed on the floor. We also confirmed from video footage that participants’ arms remained held tightly close to their torso for the entire trial.

Filtered and interpolated kinematic data were exported for further processing using MATLAB (R2014a, The MathWorks, Inc.). Whole body COM was calculated using a 5-segement model (head-arms-trunk segment, left and right thigh segments, and left and right shank-foot segments) from previously published segment COM data (Winter *et al.*, 1998). Mediolateral extrapolated COM (X_COM_) was calculated using Equation 1.

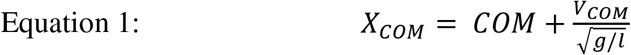

In Equation 1, *COM* is the medio-lateral position of the COM, *V*_*COM*_ is the medio-lateral velocity of the COM, *g* is the gravity constant (9.81 m/s^2^), and *l* is the length of the pendulum (mean length of left and right legs multiplied by 1.34).^2^ Leg length was the vertical distance between greater trochanter and lateral malleolus markers during the quiet standing calibration trial.

MOS was calculated relative to the medial (MOS_M_) and lateral (MOS_L_) borders of the BOS; the BOS was defined by the borders of the beam. For left steps (right stance) Equation 2 and 3 were used to calculate the MOS.

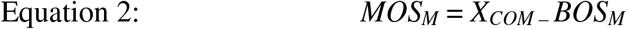

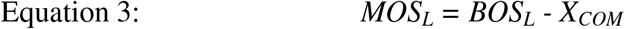

The distance of the COM was also calculated relative to the medial (COM_M_) and lateral (COM_L_) boundaries of the BOS (i.e., borders of balance beam) using Equations 4 and 5 for left steps (right stance).

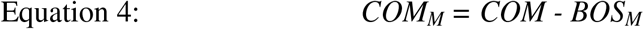

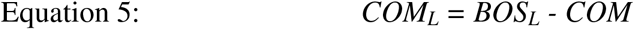

For right steps (left stance) the same equations were used (Equations 2–5) but MOS_M_, MOS_L_, COM_M_ and COM_L_ were multiplied by −1; thus, values were positive when X_COM_ or COM were inside the beam and negative when outside the beam (see also Figure 1). The minimum mediolateral MOS (minMOS) and COM (minCOM) were the lowest values of MOS_M_ or MOS_L_, and COM_M_ or COM_L_, respectively, during each step. For negative values of minMOS and minCOM, we determined the direction of ‘instability’ by determining if the medial or lateral value was first negative during the step. The following variables were also extracted: the time when MOS and COM first became negative, relative to the start and of the step; forward progression speed (mean antero-posterior velocity of the COM during the step); and step duration.

**Figure 1:**
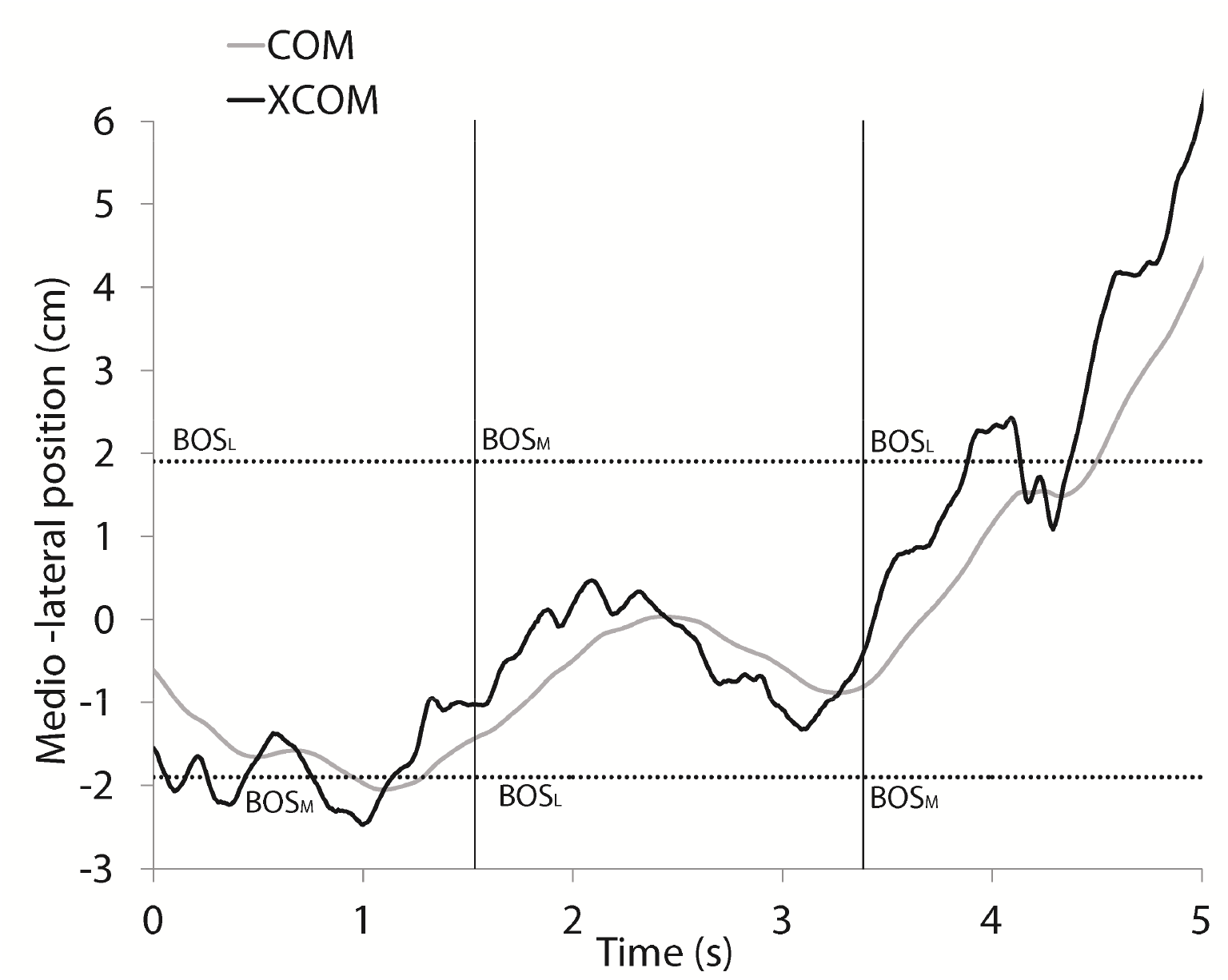
Centre of mass (COM) position and extrapolated centre of mass (XCOM). Data are shown over three consecutive steps. Each step is separated by a vertical line, and the boundaries of the beam are indicated by the dotted horizontal lines. For the first step (left swing, right stance), both the COM and XCOM cross the beam boundary (i.e., the medial border of the base of support; BOS_M_), but the step was successful. For the second step (right swing, left stance), both the COM and XCOM were within the boundaries of the beam and the step was successful. For the third step (left swing, right stance), the XCOM and COM cross the boundary of the beam (i.e., the lateral border of the base of support; BOS_L_), and the step was failed.

### Statistical analysis

Steps were categorized as those with positive and negative minMOS/minCOM, and successful/failed steps. We then calculated the number of steps where the minMOS and minCOM correctly predicted successful or failed steps. When minMOS or minCOM were negative in the lateral direction, a crossover step (i.e., step landing lateral to the stance limb) would be required to regain stability. Crossover stepping was observed infrequently and, therefore, is likely not an ideal strategy to regain stability for this particular balance beam task. Instead, participants seemed to place the swing limb quickly on the beam and regained stability in the subsequent step by stepping laterally with the stance limb. Thus, the following situations would constitute a correct prediction: 1) positive minMOS/minCOM during a successful step; 2) negative minMOS/minCOM prior to a failed step; 3) or negative minMOS/minCOM in the lateral direction during a successful step, followed immediately by failed step. Chi-square test was used to compare the proportion of correct predictions for minMOS to minCOM.

We conducted additional analysis of step characteristics to help explain errors in predicting successful or failed steps. For steps that had a negative minMOS or minCOM in the medial direction, repeated measures analysis of variance (ANOVA) was used to compare the timing of negative minMOS or minCOM, step duration, and speed of progression between successful and failed steps. Only participants who had both failed and successful steps with negative minMOS and/or minCOM in the medial direction were included in this analysis. Additionally, for successful steps with negative minMOS or minCOM, we compared the magnitude of minMOS or minCOM, timing of negative minMOS or minCOM, step duration and speed of progression between steps where the subsequent step was successful or failed.

## RESULTS

One trial was excluded from analysis for each of four participants due to a high number of obscured markers. Altogether, participants took a total of 239 steps during the trials, 141 of which were successful and 98 of which were failed.

### Accuracy of measures in predicting step outcome

There were 5 steps with positive minMOS, and all of these were successful (Figure 2). There were 234 steps with negative minMOS. For 92 steps, minMOS was negative in the lateral direction; 84 of these were successful, with the subsequent step being failed for 37 and successful for 47 steps. The majority of steps where minMOS was negative in the medial direction were failed (90/144; 62.5%).

**Figure 2:**
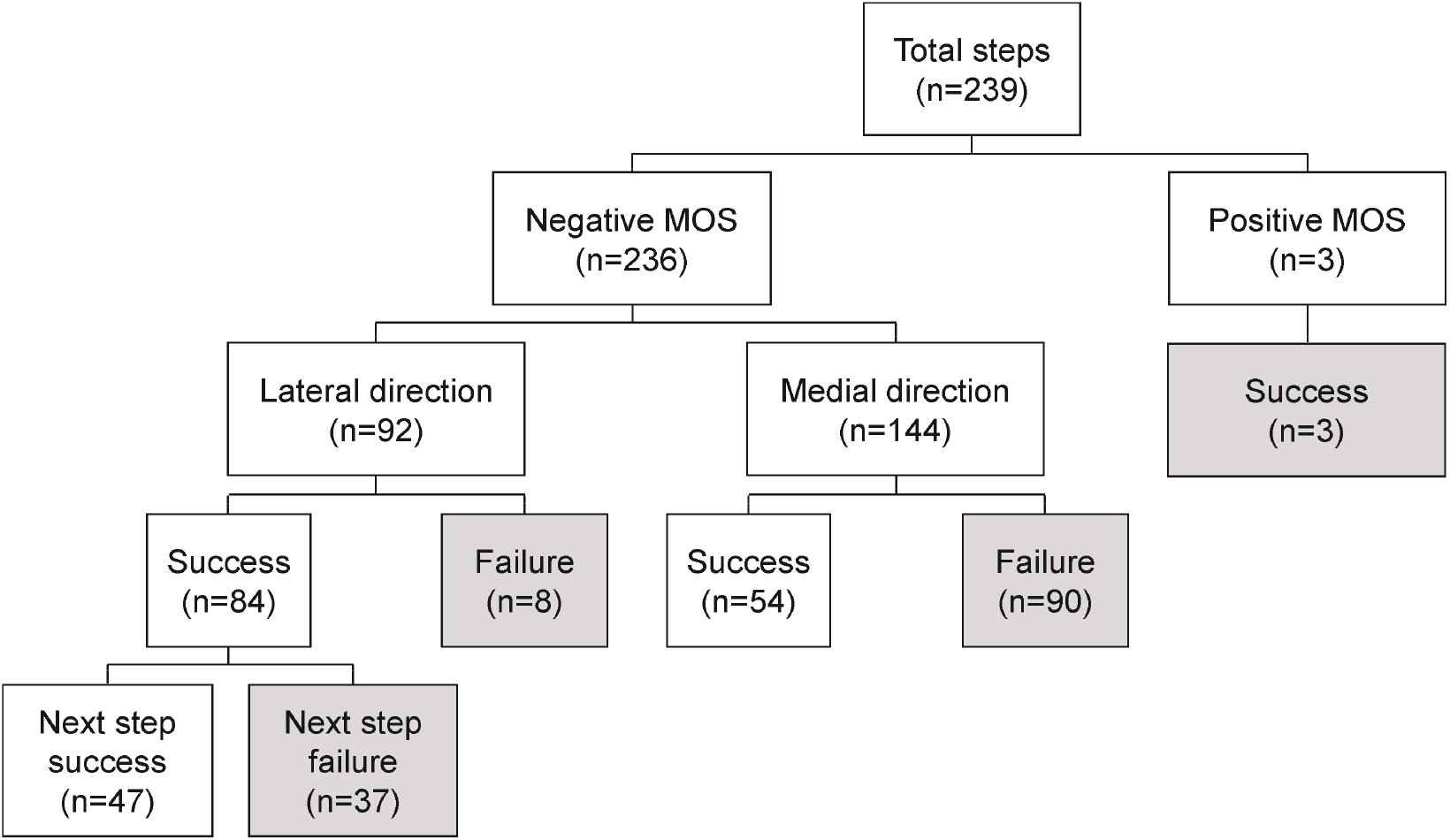
Ability of MOS to predict successful or failed steps. For the 239 beam-walking steps, 234 had a negative minMOS and 5 had a positive minMOS. Boxes with grey shading indicate that the minMOS correctly predicted the outcome of the step (success or failure).

All 32 steps with positive minCOM were successful (Figure 3). There were 207 steps with negative minCOM. Of the 73 steps with negative minCOM in the lateral direction, 67 (96%) were successful, with the subsequent step being failed 35 times, and successful for 32 steps. The majority of steps with negative minCOM in the medial direction failed (92/134; 68.7%).

**Figure 3:**
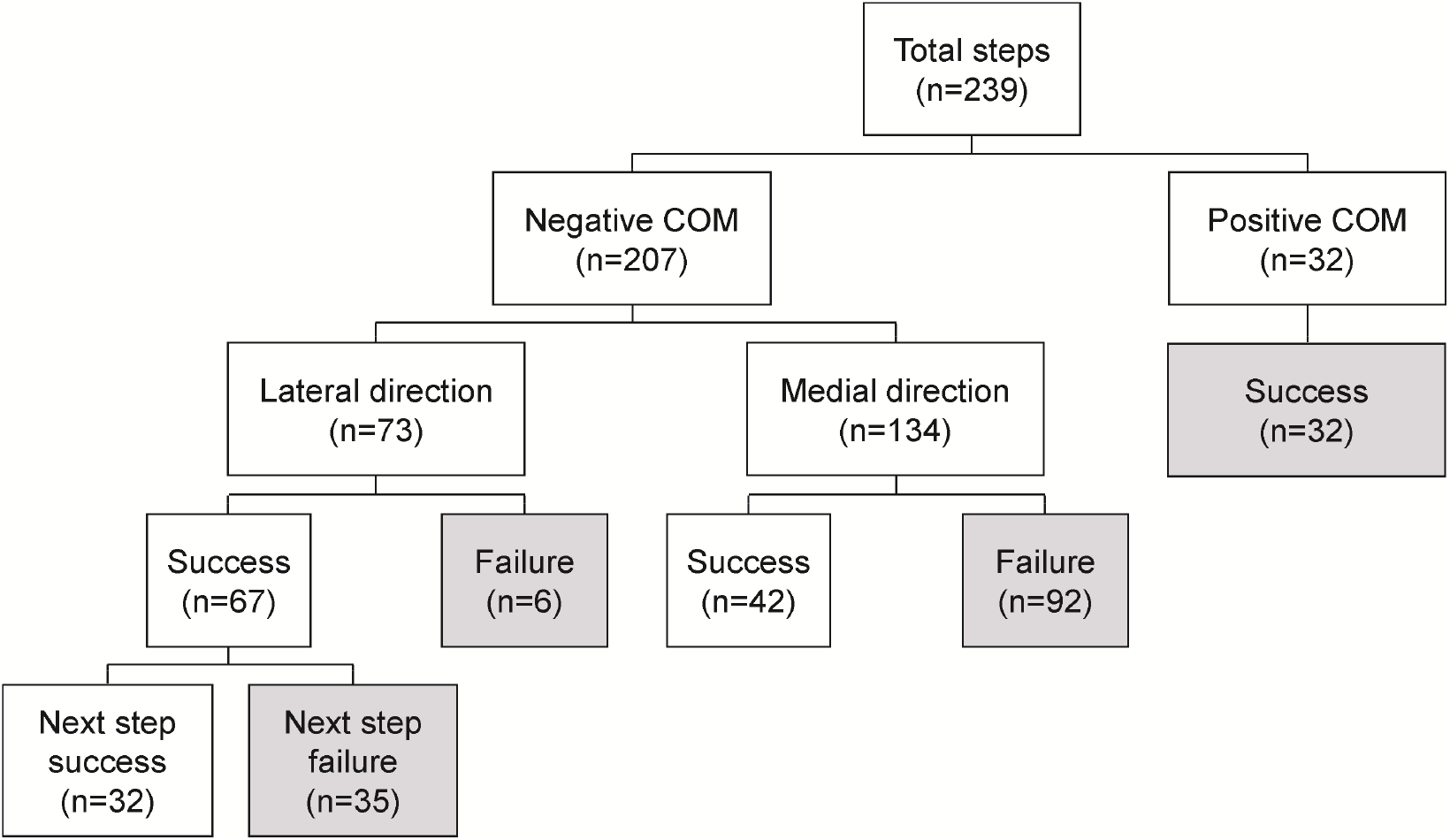
Ability of COM position to predict successful or failed steps. For the 239 beam-walking steps, 207 had a negative minCOM and 32 had a positive minCOM. Boxes with grey shading indicate that the minCOM correctly predicted the outcome of the step (success or failure).

The minMOS correctly predicted step outcome for 57.7% of steps (138/239), whereas minCOM correctly predicted step outcome for 69.0% of steps (165/239). This difference in prediction accuracy was statistically significant (χ^2^=6.57; p=0.010).

### Comparison of step characteristics between successful and failed steps

One participant was excluded from the comparison of step characteristics between steps with negative and positive minCOM as they did not have both successful and failed steps with a negative minCOM. For steps with a negative minMOS or minCOM in the medial direction, failed steps tended to have quicker step time (minMOS: F_1,8_=29.53, p=0.0006; minCOM: F_1,7_=34.75, p=0.0006), but had slower speed of progression than successful steps (minMOS: F_1,8_=16.56, p=0.0036; minCOM: F_1,7_=14.83, p=0.0063; Table 1). For steps with negative minMOS or minCOM, there was no significant difference between failed and successful steps in timing of negative minMOS or minCOM, relative to the start of the step (minMOS: F_1,8_=2.48, p=0.15; minCOM: F_1,7_=2.02, p=0.20).

**Table 1:** Comparison of step characteristics between successful and failed steps. Values shown are least square means with 95% confidence interval in brackets. The p-value is for the repeated measures ANOVA comparing successful and failed steps. Only trials with negative minMOS or negative minCOM in the medial direction are included. One participant was excluded from the minCOM analysis as they did not have both successful and failed steps with a negative minCOM.

|  | Failed step | Successful step | p-value |
| --- | --- | --- | --- |
| **minMOS** |  |  |  |
| Number of steps | 90 | 54 |  |
| Time of negative minMOS (ms) | 90 [29, 150] | 221 [140, 303] | 0.15 |
| Step duration (ms) | 983 [858, 1107] | 1514 [1347, 1682] | 0.0006 |
| Speed of progression (m/s) | 0.15 [0.14, 0.17] | 0.22 [0.19, 0.24] | 0.0036 |
| **minCOM** |  |  |  |
| Number of steps | 74 | 42 |  |
| Time of negative minCOM (ms) | 193 [78, 308] | 339 [185, 493] | 0.20 |
| Step duration (ms) | 983 [829, 1136] | 1594 [1389, 1799] | 0.0006 |
| Speed of progression (m/s) | 0.15 [0.13, 0.17] | 0.21 [0.18, 0.24] | 0.0063 |

For successful steps with negative minMOS or minCOM, the minMOS or minCOM had a significantly larger negative value when the next step was failed compared to when the next step was successful (F_1,8_>10.07, p<0.014; Table 2). There were no significant differences in timing of negative minMOS/minCOM, step time, or speed of progression, between subsequent failed and successful steps for successful steps with negative minMOS/minCOM (F_1,8_<3.15, p>0.11).

**Table 2:**
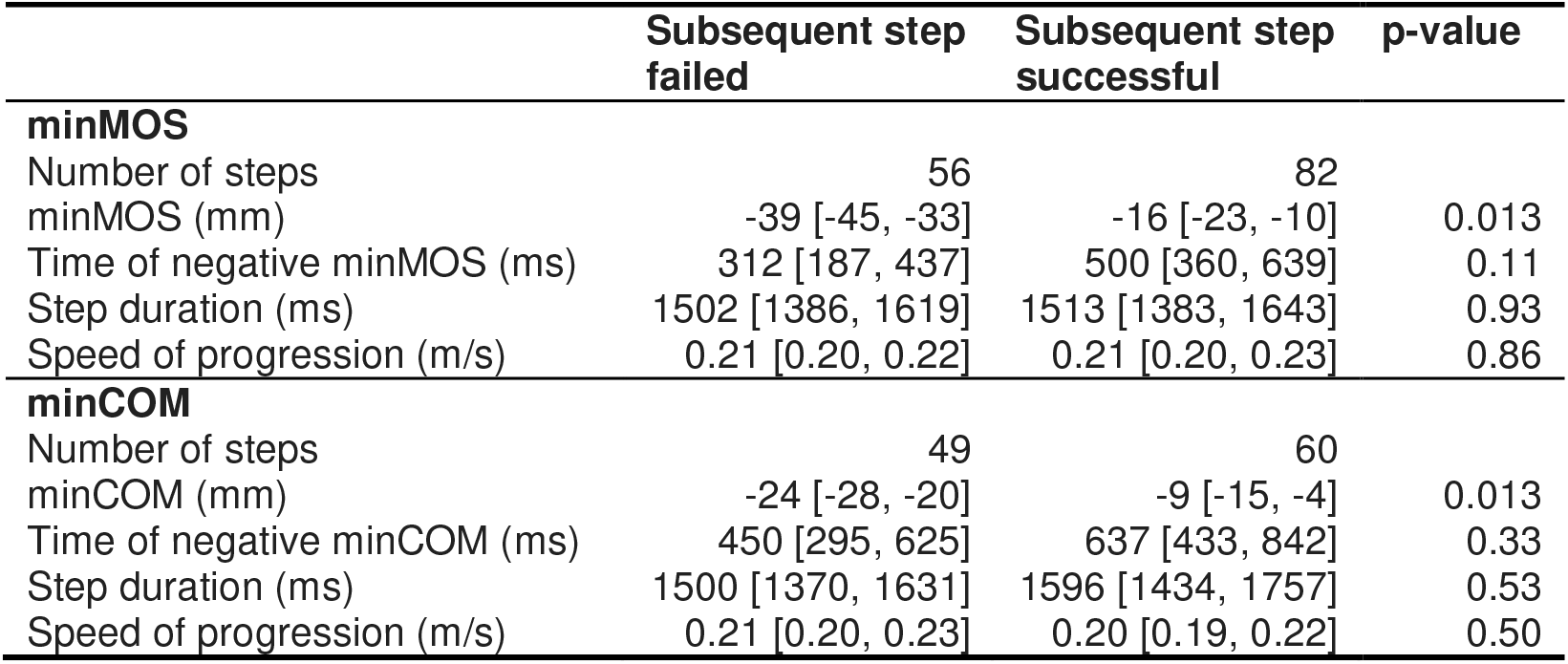
Comparison of step characteristics between subsequent failed and successful steps. Values shown are least square means with 95% confidence intervals in brackets. The p-value is for the repeated measures ANOVA comparing successful and failed steps. Only successful steps with negative minMOS or minCOM were included in this analysis.

## DISCUSSION

The purpose of this study was to determine the validity of mediolateral MOS and COM position for describing stability in the context of beam-walking. During a beam-walking task, the BOS has a fixed, constant width in the mediolateral direction. We hypothesized that the minMOS and minCOM should remain positive throughout successful steps on the beam, and instances of negative minMOS or minCOM would result in ‘instability’ and necessitate stepping off the beam (i.e., a failed step).

In agreement with our hypothesis, all steps with positive minMOS and minCOM were successful. However, contrary to our hypothesis, a large number of successful steps had a negative minMOS or a negative minCOM. Speed of progression was faster for successful steps with negative minMOS and/or minCOM than failed steps. Previous work has shown increasing step frequency can improve mediolateral MOS stability (Hak *et al.*, 2013). Thus in our study, while participants were instructed to walk to the sound of a metronome (and thus step frequency was controlled), participants may have walked faster to maintain mediolateral stability despite the negative minMOS or minCOM. However it is unclear to what extent participants directed effort towards completing the goal of the task (successfully stepping on beam to metronome) versus maintaining balance and avoiding a fall.

When the XCOM or COM crossed the lateral boundary of the beam, a lateral ‘crossover’ step with the swing limb would be required to regain stability. We only observed three of these crossover steps and assumed that, in these cases of lateral instability, the participants’ strategy would be to step on the beam quickly with the swing limb (thus successfully stepping based on our parameters), and then step off the beam with the stance limb. In which case, successful steps that were followed by an unsuccessful step may have all been part of the same overall “failure” process. This is partially supported by our findings, in that successful steps with a negative minMOS or minCOM were faster than a failed step, indicating that participants may have tried to maintain the goal of the task (staying atop the beam) while in the overall act of failing.

Not all successful steps with a negative minMOS or minCOM were immediately followed by a failed step. Our findings indicate examining the task scenario as inside/outside beam boundary may be over-simplified, in that participants may be able to use other strategies to prevent falling from the beam when negative minMOS and minCOM values are relatively small. Previous investigators have demonstrated that negative MOS values can be compensated for through major trunk movements (Hof *et al.*, 2005). This may have occurred in our task, as trunk movements are used more often when the BOS is very narrow (Otten, 1999; Hof *et al.*, 2007). Additionally, ankle musculature has been shown to contribute to mediolateral balance when the feet are in tandem and thus the width of the BOS is small (Winter *et al.*, 1996). Both of these strategies may have been used by participants to help complete multiple successful steps with negative minMOS or minCOM, as our findings demonstrated a significant difference in minMOS and minCOM between successful and failed steps following a successful step with negative minMOS or minCOM. Thus, a number of potential strategies could have been implemented by participants to complete successful steps while minMOS or minCOM were negative.

There are some limitations in our study. Firstly, a number of assumptions exist both when computing our COM model, and when computing the MOS. Both of these model assumptions may have played a role in our findings of small, negative minMOS or minCOM values existing during successful stepping. This may be especially true of the MOS the inverted pendulum model of the XCOM may not accurately describes mediolateral stability during a narrow beam walking task. Secondly, we cannot definitively determine what level of negative minMOS or minCOM was tolerable for participants to maintain successful stepping with alternative strategies. While our findings indicated that participants were able to complete a successful step with a minMOS of approximately −16mm and a minCOM of approximately −9mm, it is unclear if this is the mechanical limit of stability when factoring in alternate strategies, or the ‘functional’ limit of stability, as perceived by the central nervous system via somatosensory feedback from the bottom of the feet (Horak, 2006). Previous work has shown that individuals have a functional base of support, which can change or decrease due to age or clinical dysfunction (King *et al.*, 1994; Wolfson, 2001). Thus, we cannot for certain determine whether our findings represent the mechanical or functional limits of stability for this balance beam walking task.

## CONCLUSION

Previous authors have suggested that an individual is unstable when the COM (Shumway-Cook and Woollacott, 2007) or XCOM (Hof *et al.*, 2005) is outside the BOS. The results of this study suggest that classifying the system as stable or unstable based on whether the COM or XCOM is inside or outside the BOS may be over-simplistic; this classification correctly predicted stability during the beam-walking task in only 69% and 58% of cases, respectively. However, the magnitude of excursion of the COM or XCOM outside of the BOS was related to stability; while participants could tolerate a small excursion of the COM or XCOM outside of the BOS, a larger excursion was more likely to lead to instability. Thus, simply classifying of the system as stable or unstable based a strict threshold of whether the COM or XCOM is inside or outside the BOS may not be valid. Alternatively, treating the COM position relative to the BOS and the MOS as continuous variables is useful for informing stability of the system.

## Acknowledgements

This study was supported by the Natural Sciences and Engineering Research Council of Canada (RGPIN-2014-04199). We also acknowledge the support of the Toronto Rehabilitation Institute; equipment and space have been funded with grants from the Canada Foundation for Innovation, Ontario Innovation Trust, and the Ministry of Research and Innovation. AM holds a New Investigator Award from the Canadian Institutes of Health Research (MSH-141983). AHH was supported by a Trainee Award from the Heart and Stroke Foundation Canadian Partnership for Stroke Recovery. The funders had no role in the collection, analysis, and interpretation of data; in the writing of the manuscript; and in the decision to submit the manuscript for publication. We would also like to thank Ming Ma, Rebecca Graham, Patrick Antonio, and Shajicaa Sivakumaran for their help with data collection and processing.

## REFERENCES

Bruijn, S. M., Meijer, O. G., Beek, P. J., van Dieën, J. H., 2013. Assessing the stability of human locomotion: a review of current measures. Journal of The Royal Society, Interface 10, 20120999.

Bruijn, S. M., van Dieën, J. H., 2018. Control of human gait stability through foot placement. 15, 20170816.

Ghoussayni, S., Stevens, C., Durham, S., Ewins, D., 2004. Assessment and validation of a simple automated method for the detection of gait events and intervals. 20, 266–272.

Hak, L., Houdijk, H., Beek, P. J., Van Dieën, J. H., 2013. Steps to take to enhance gait stability: the effect of stride frequency, stride length, and walking speed on local dynamic stability and margins of stability. 8, e82842.

Hof, A. L., 2008. The ‘extrapolated center of mass’ concept suggests a simple control of balance in walking. Human Movement Science 27, 112–125.

Hof, A. L., Gazendam, M. G. J., Sinke, W. E., 2005. The condition for dynamic stability. Journal of Biomechanics 38, 1–8.

Hof, A. L., van Bockel, R. M., Schoppen, T., Postema, K., 2007. Control of lateral balance in walking. Experimental findings in normal subjects and above-knee amputees. 25, 250–258.

Horak, F. B., 2006. Postural orientation and equilibrium: what do we need to know about neural control of balance to prevent falls? 35, ii7–ii11.

King, M. B., Judge, J. O., Wolfson, L., 1994. Functional base of support decreases with age. 49, M258–263.

Otten, E., 1999. Balancing on a narrow ridge: biomechanics and control. 354, 869–875.

Pai, Y.-C., Patton, J., 1997. Center of mass velocity-position for balance control. Journal of Biomechanics 30, 347–354.

Shumway-Cook, A., Woollacott, M. H., 2007. Motor control: translating research into clinical practice. Philadelphia, Lippincott Williams & Wilkins.

Sivakumaran, S., Schinkel-Ivy, A., Masani, K., Mansfield, A., 2018. Relationship between margin of stability and deviations in spatiotemporal gait features in healthy young adults. 57, 366–373.

Winter, D. A., 1995. Human balance and posture control during standing and walking. 3, 193–214.

Winter, D. A., Patla, A. E., Prince, F., Ishac, M., Gielo-Perczak, K., 1998. Stiffness control of balance in quiet standing. 80, 1211–1221.

Winter, D. A., Prince, F., Frank, J. S., Powell, C., Zabjek, K. F., 1996. Unified theory regarding A/P and M/L balance in quiet stance. 75, 2334–2343.

Wolfson, L., 2001. Gait and balance dysfunction: a model of the interaction of age and disease. 7, 178–183.

